# A method for authenticating the fidelity of *Cryptococcus neoformans* knockout collections

**DOI:** 10.1101/2024.12.03.626716

**Authors:** Ella Jacobs, Quigly Dragotakes, Madhura Kulkarni, Amanda Dziedzic, Anne Jedlicka, J. Marie Hardwick, Arturo Casadevall

## Abstract

Gene knockout strain collections are important tools for discovery in microbiology. *Cryptococcus neoformans* is the only human pathogenic fungus with an available genome-wide deletion collection, and this resource is widely used by the research community. We uncovered mix-ups in the assembly of the commercially available *C. neoformans* deletion collection of ∼6,000 unique strains acquired by our lab. While pursuing the characterization of a gene-of-interest, the corresponding deletion strain from the *C. neoformans* KO collection displayed several interesting phenotypes associated with virulence. However, RNAseq analysis identified transcripts for the putative knockout gene, and the absence of transcripts for a different knockout strain found in the same plate position in an earlier partial knockout collection, raising the possibility that plates from one collection were substituted for the other. This was supported by determining the size of the nourseothricin (NAT)-resistance cassette used to generate the two separate knockout libraries and was confirmed by RNAseq and genome sequencing. Here we report that our KN99 collection is comprised of mixed plates from two independent KO libraries and present a simple authentication method that other investigators can use to distinguish the identities of these KO collections.

**Importance:** Gene knockout strain collections are important tools for discovery in microbiology. *Cryptococcus neoformans* is the only human pathogenic fungus with an available genome-wide deletion collection, and this resource is widely used by the research community. Here we report that our KN99 collection is comprised of mixed plates from two independent KO libraries and present a simple authentication method that other investigators can use to distinguish the identities of these KO collections. Above all, this article serves as a reminder to 2015 library collection strains before undertaking large phenotyping experiments.

## Introduction

In the field of microbiology, genome-wide gene deletion collections are important investigative tools to rapidly screen for genes associated with specific phenotypes. For studies of *Cryptococcus neoformans*, several gene deletion collections are available [1, 2], including the genome-wide knockout collection containing ∼4,700 strains generated in the Madhani laboratory, an arduous feat compared to the more tractable model yeast *Saccharomyces cerevisiae* [3–5].

The first Madhani *C. neoformans* gene knockout (KO) collection was generated in the H99W background strain (2008 collection of ∼1300 KO strains) [6]. H99W, also known as “Wimp”, is a laboratory adapted strain with attenuated virulence and melanization defects attributed to a 7 bp insertion in the *LMP1* gene [7, 8]. Given the severe virulence attenuation of the H99W strain, a subsequent genome-wide deletion collection was generated in the virulent KN99α background strain, arrayed in 96-well plates and distributed in three phases, designated as 2015 (22 plates), 2016 (20 plates) and 2020 (7 plates) collections containing one KO strain per well [9]. Knockout strains were generated by gene replacement/disruption via homology directed repair using gene-specific ∼1 kb homology arms flanking a nourseothricin (NAT) cassette [9].

Our 2015 and 2016 KO collections were purchased from the Fungal Genetics Stock Center (FGSC) in March 2018 and the 2020 KO collection in March 2021. These purchased collections have proved useful for screening and identifying genes associated with non-lytic exocytosis [10] protein secretion [11], and cell death [12]. However, during the characterization of several deletion strains selected to follow up on proteomics studies, we realized that many strains from our 2015 collection did not contain the expected gene disruption, despite being NAT resistant.

We tested plates of our 2015 collection and found that rather than individual well mix-ups or well-well contamination, whole plates from the KN99 collection appear to have been swapped with 2008 collection plates. The 2008 KO collection contains strains are arrayed in 14 96-well plates. Based on the strains we tested, only plates within our first 14 plates of the 2015 KO collection were impacted. Overall, a total of nine non-consecutive plates from the 2008 collection appeared to be directly swapped into the 2015 collection, while no mix-ups were detected for the strains tested from the 2016 and 2020 collections. We developed a simple, cost-effective method of testing deletion strains purchased from the Fungal Genetics Stock Center (FGSC) by comparing NAT cassette sizes. However, locus-specific validation is required to fully confirm plate integrity. This protocol will allow screening to quickly assess potential fidelity issues for follow-up.

## Results

### Generation of primers to reveal the location and prescence of NAT cassettes within our 2015 KO collection strains

Using overlap PCR, the Madhani lab constructed deletion cassettes containing the NAT-resistance gene plus 1 kb flanking homology arms. These cassettes were subsequently transformed and inserted into the *C. neoformans* gene of interest by homology directed repair (HDR) [9]. The primers used by the Madhani lab to generate the 2015 KO collection are available on the FGSC website (Fig 1A). Primer pairs W1+W2 and W5+W6 amplify the flanking 5’ and 3’ recombination arms, respectively. Primer pairs W3 + W4 (not included in the available primer dataset) were used to amplify the NAT resistance cassette, which was fused to the genomic recombination arms (Fig 1A).

**Figure 1.**
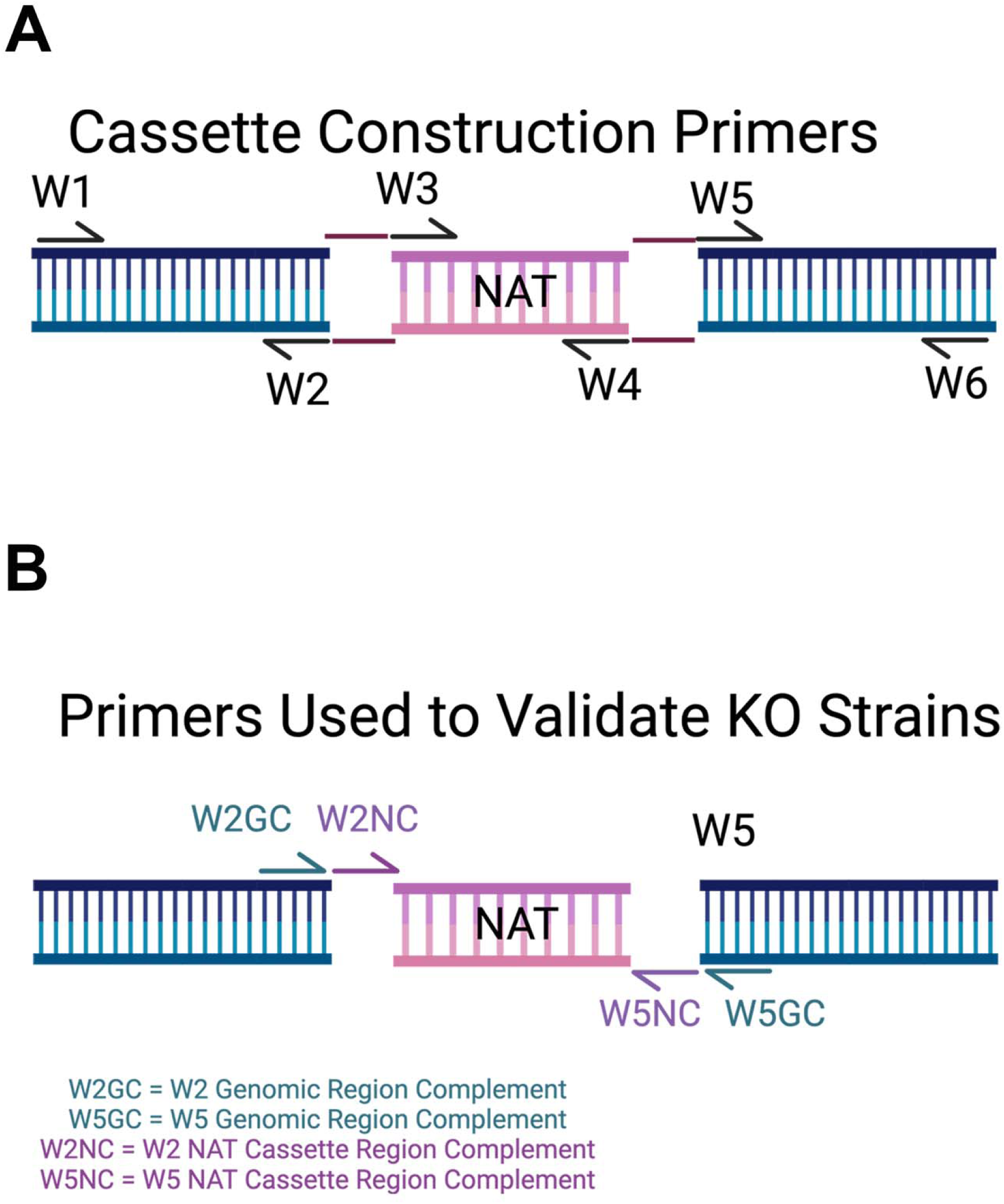
Primer design schematics for KO construction and validation **A)** Deletion cassette construction adapted from [9]: W1, W2, W5, and W6 primers are publicly available for each strain on the FGSC website. W3 and W4 primers are not publicly available. Created with BioRender. **B)** Strain verification schematic. The reverse complement of the genomic region of W2 (W2GC) and W5 (W5GC) were used to confirm deletion fidelity at each locus.

To amplify the NAT cassette in any KO strain, regardless of location, we generated primers complement of the 5’ of W2 + W5 NAT cassette region, designated as W2 NAT Complement (W2NC) and W5 NAT Complement (W5NC) (Figure 2A Table 1). According to the FGSC spreadsheet, the 5’ end of the W2 + W5 primer pairs is the same across all 2015 KO constructs. Therefore, this primer pair could be used on any deletion strain to amplify the NAT cassette as it contains no locus-specific base pairs.

**Figure 2:**
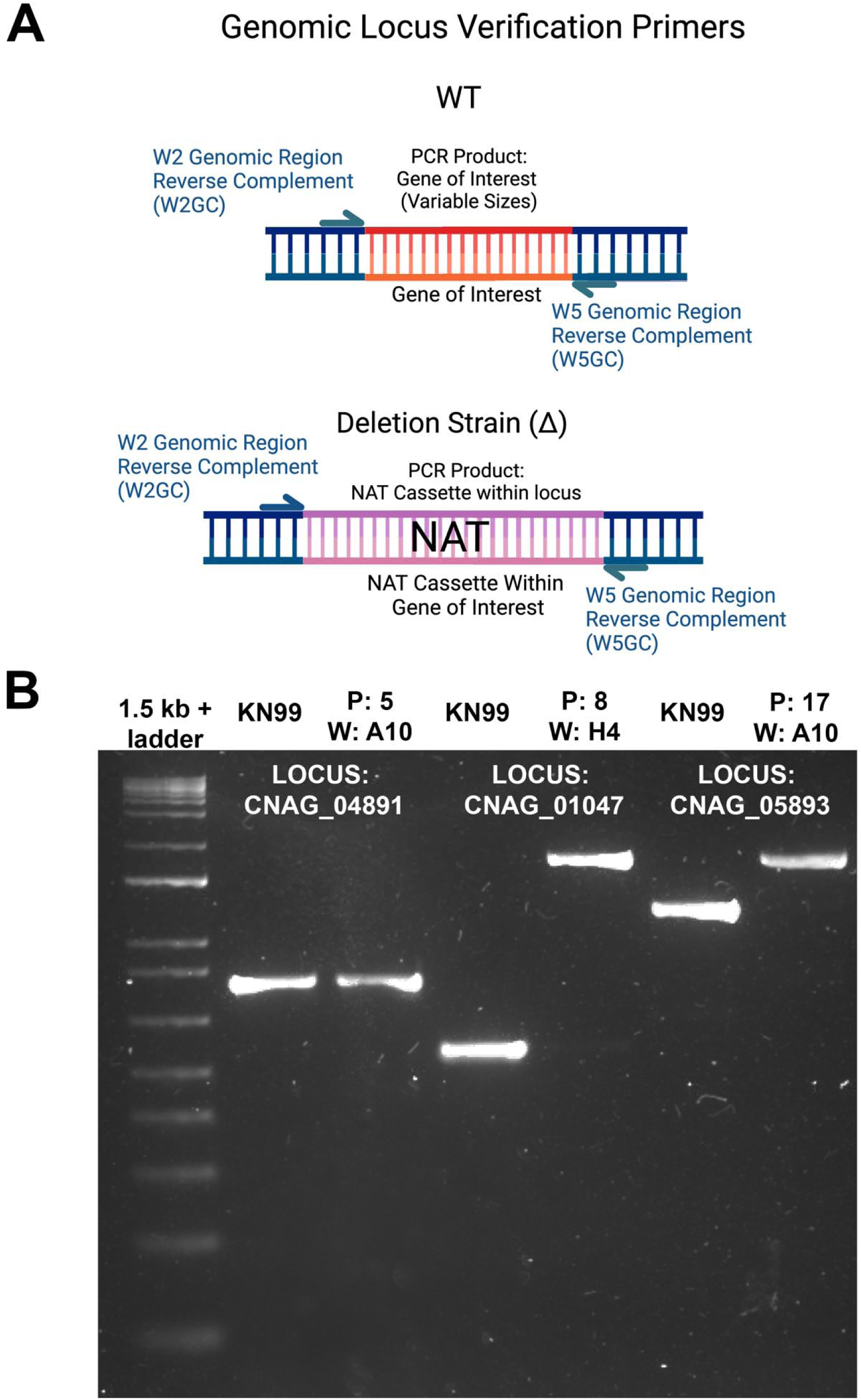
PCR validation of KO collection strains targeting the genomic locus **A)** In WT KN99, these primer pairs amplify the gene of interest. In KO strains, these primer pairs will only amplify the NAT cassette. **B)** Example genomic locus strain verification 1.5% agarose gel using the methodology in panel B. Primers targeted the reverse complement of the genomic locus specific sequence in the W2 (W2GC) and W5 (W5GC). K-denoted lanes contain WT (KN99) PCR products and Δ-denoted lanes contain putative deletion strain PCR products. Numbers refer to the CNAG locus corresponding to the W2 W5 sequence location. CNAG_04891Δ did not contain a NAT cassette within CNAG_04891, as the WT and deletion strain PCR products are the same size. Both CNAG_01047Δ and CNAG_05893Δ contain NAT cassettes in the expected locus.

**Table 1:**
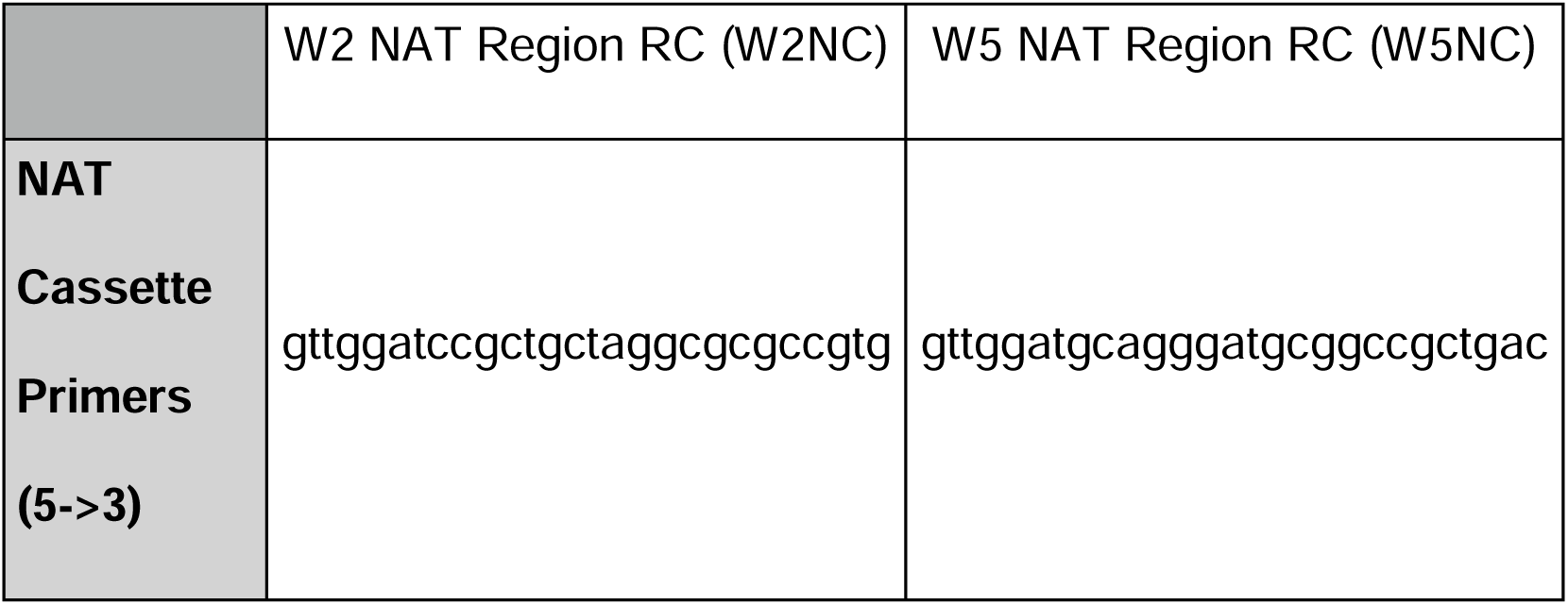
NAT cassette PCR primers.

We generated two types of primers to verify either the location or presence of the NAT drug selection marker in our copy of the KO collections (Fig 1B). To verify the location of the NAT cassette in the correct KO gene locus we generated primers that bind the genome immediately flanking the NAT cassette. These primers were adapted from W2 and W5 sequences, complementary to the 3’ 27-nucleotides of primers W2 and W5, indicated as W2 Genomic Complement (W2GC) and W5 Genomic Complement (W5GC) (Fig 1B). The sequence of W2GC and W5GC differs for each gene (Table 2).

**Table 2:**
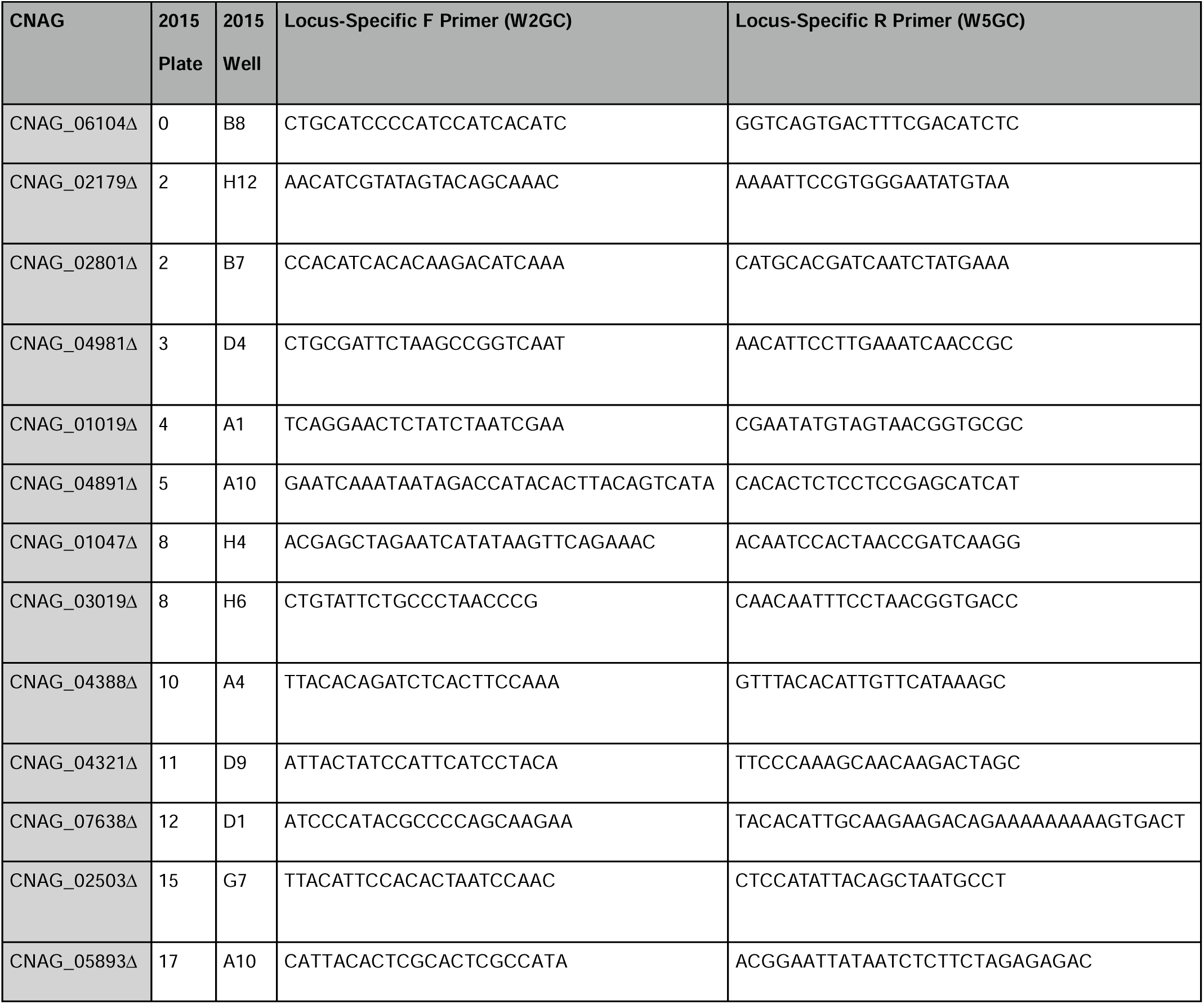
W2GC and W5GC locus-specific primers.

### Targeting genomic loci of our 2015 KO collection library strains via PCR

W3GC and W4GC primer sets for 13 of genes were used to amplify genomic DNA isolated from both WT and KO strains in the 2015 collection (Fig 2A), and the sizes of amplified products were compared on agarose gels (Fig 2B, Table 3). Although initially unexpectedly, genomic PCR products of 5 out of the 12 putative KO strains were the same size at the WT gene products, for example CNAG_04891 (Fig 2D). This suggested no gene deletion of the expected gene despite being NAT-resistant. In contrast, other deletion strains from the 2015 collection appeared to contain the same size NAT cassette (∼1.7 kb), which differed from the expected sizes of their respective WT gene, for example CNAG_01047 and CNAG_05893 (Fig 1D). It is worth noting that this methodology only works if the gene of interest is different in size from the ∼1.7kb expected NAT cassette. If not, further validation with a diagnostic restriction enzyme digestion, alternative primer design or sequencing of the amplified products is necessary. We compared the size of PCR products amplified between WT and KO strains at the genomic locus (Fig 1D, Table 2).

**Table 3:**
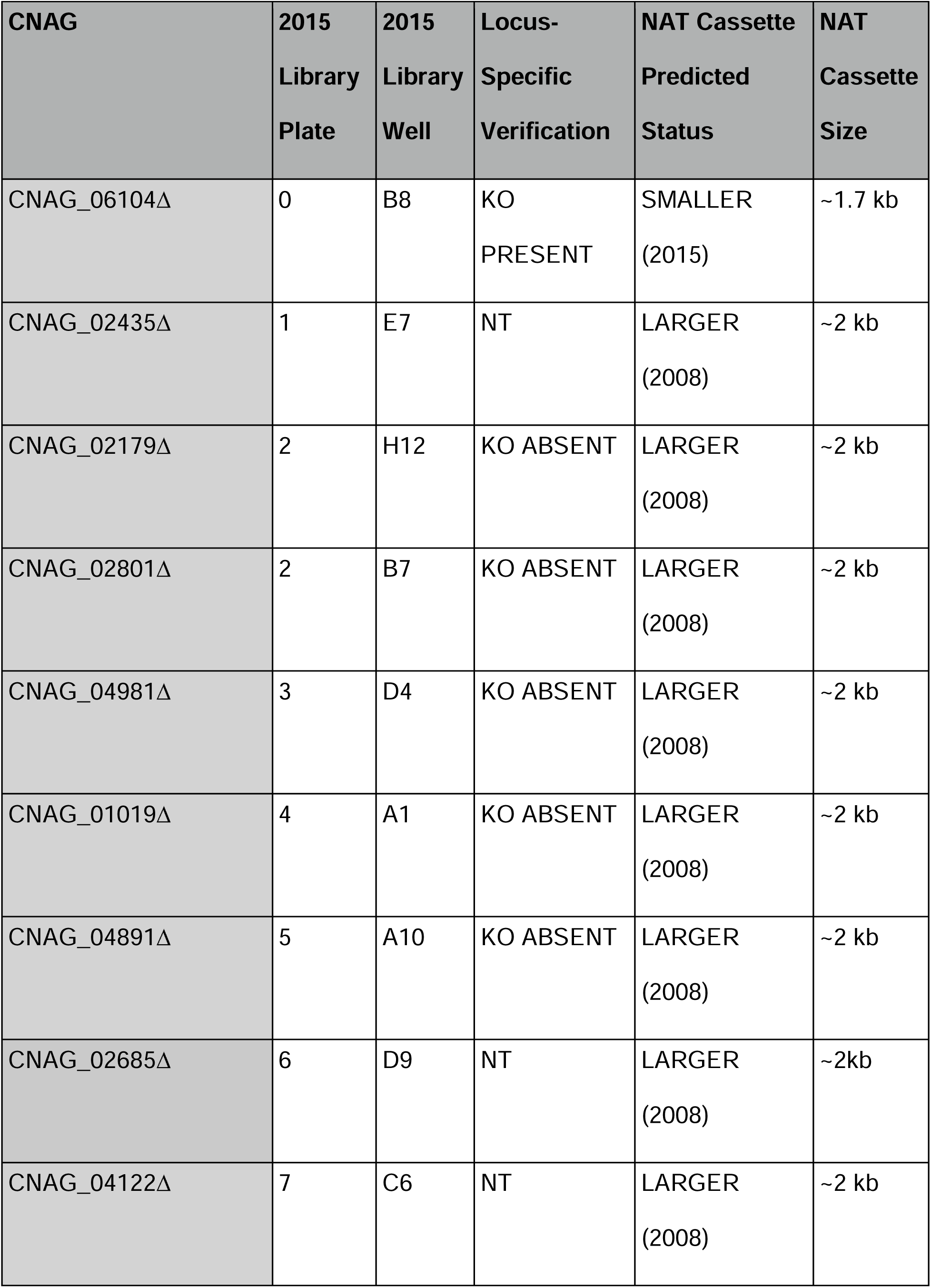
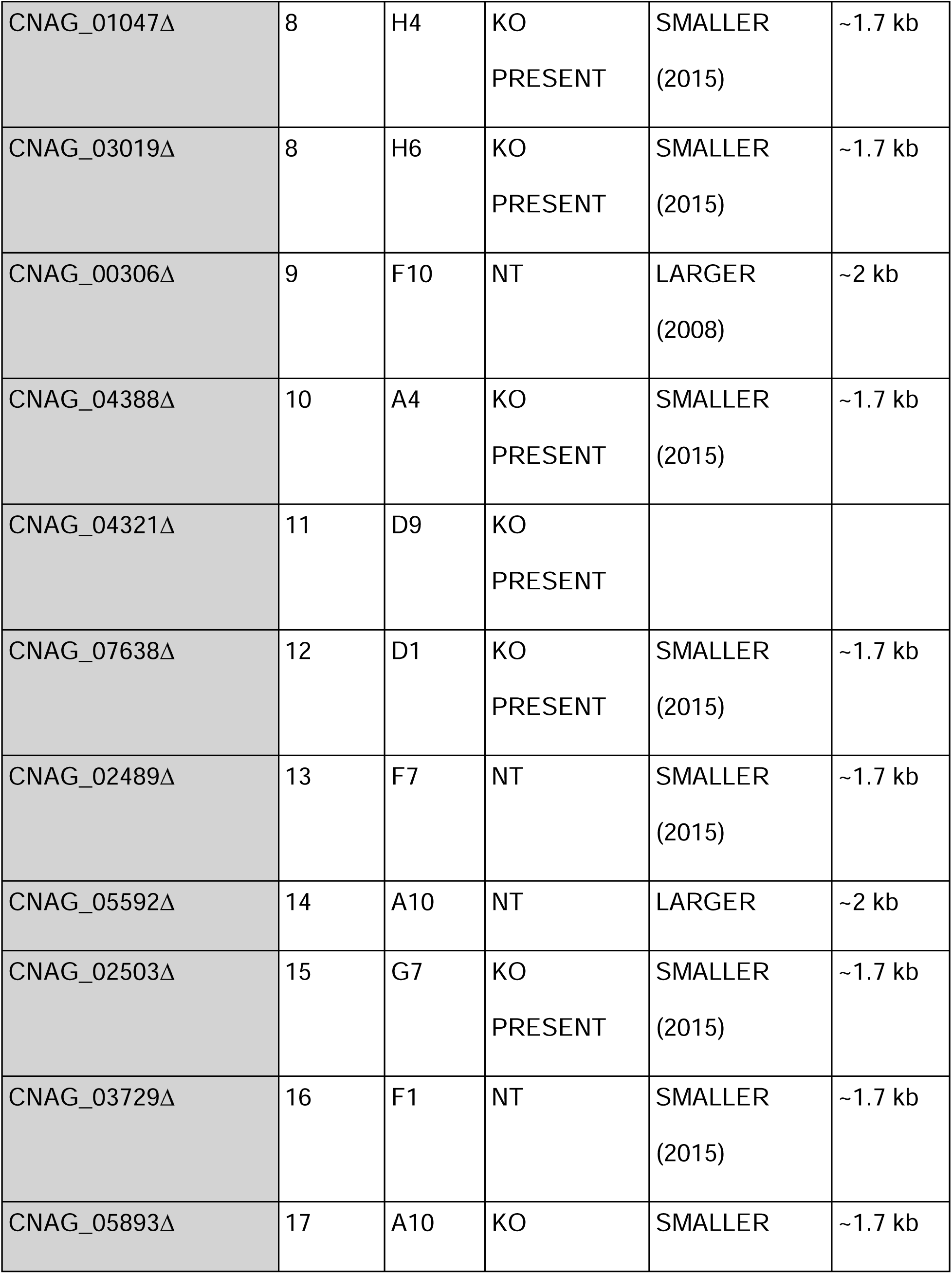
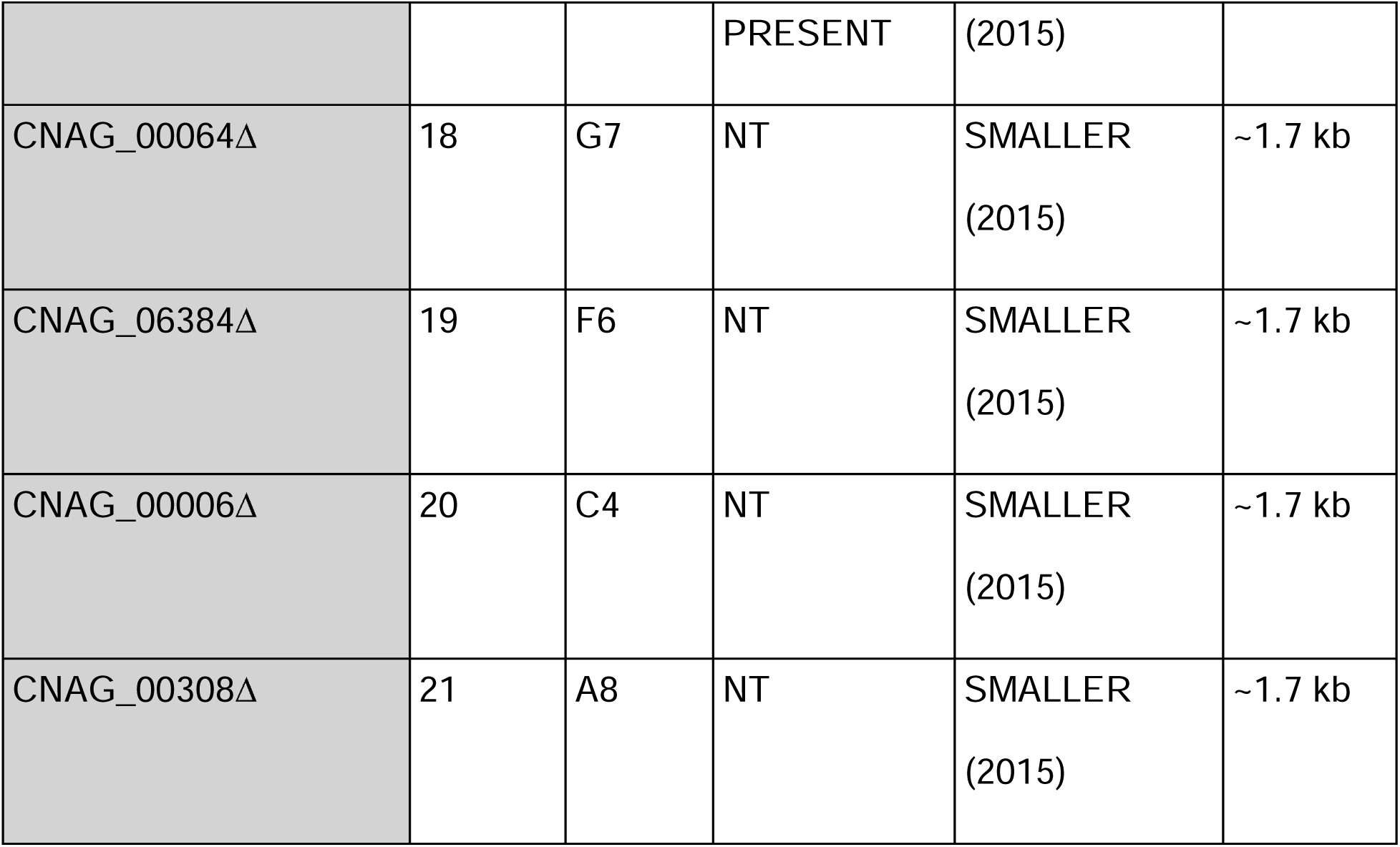
2015 KO collection strains tested using a mix of locus specific primers and NAT cassette size differences. NT indicates not tested. KO present indicates the strain contained the expected gene disruption. KO absent indicates the strain did not contain the expected disruption.

### Larger NAT cassette size correlated with strains lacking expected KO

To verify the presence of a NAT cassette in strains lacking a disruption at the expected locus, such as CNAG_04891Δ, we used the W2NC + W5NC primer pairs discussed in figure 1 B and table 1. This primer pair is capable of hybridizing to the 5’ and 3’ ends of the NAT cassette (complementary to the 5’ 22-nucleotides of primers W2 and W5), which are common to all the deletion strains (Fig 2A). As expected, no NAT cassette was amplified from WT KN99 , while all deletion strains yielded a PCR product, including strains that did not contain the expected KO (Fig 2B and 2C).

An important clue in solving the mix-up mystery became readily apparent from this analysis.

The NAT cassette amplified from strains lacking the expected KO (CNAG_01019Δ, CNAG_02179Δ, CNAG_04981Δ, CNAG_02801Δ, and CNAG_04891Δ) was ∼2 kb rather than ∼1.7 kb amplified from correct KO strains (Fig 3B. Table 2). We wondered if the differences in NAT cassette size could be due to barcoding or other potential changes that would differ between KO collections. For example, the 2008 KO collection cassettes are barcoded [6]. It appeared that the same universal primer sequences were apparently used for both 2008 and 2015 KO collections. Therefore, we conducted a systematic analysis using primers W2NC + W5NC to compare NAT cassette sizes for a representative strain from each of the 22 plates in the 2015 collection (examples in Fig 3B). In addition, while using this approach a pattern began to emerge, which suggested that the incorrect KO strains may be restricted to a subset of 96-well plates in the 2015 collection.

**Figure 3:**
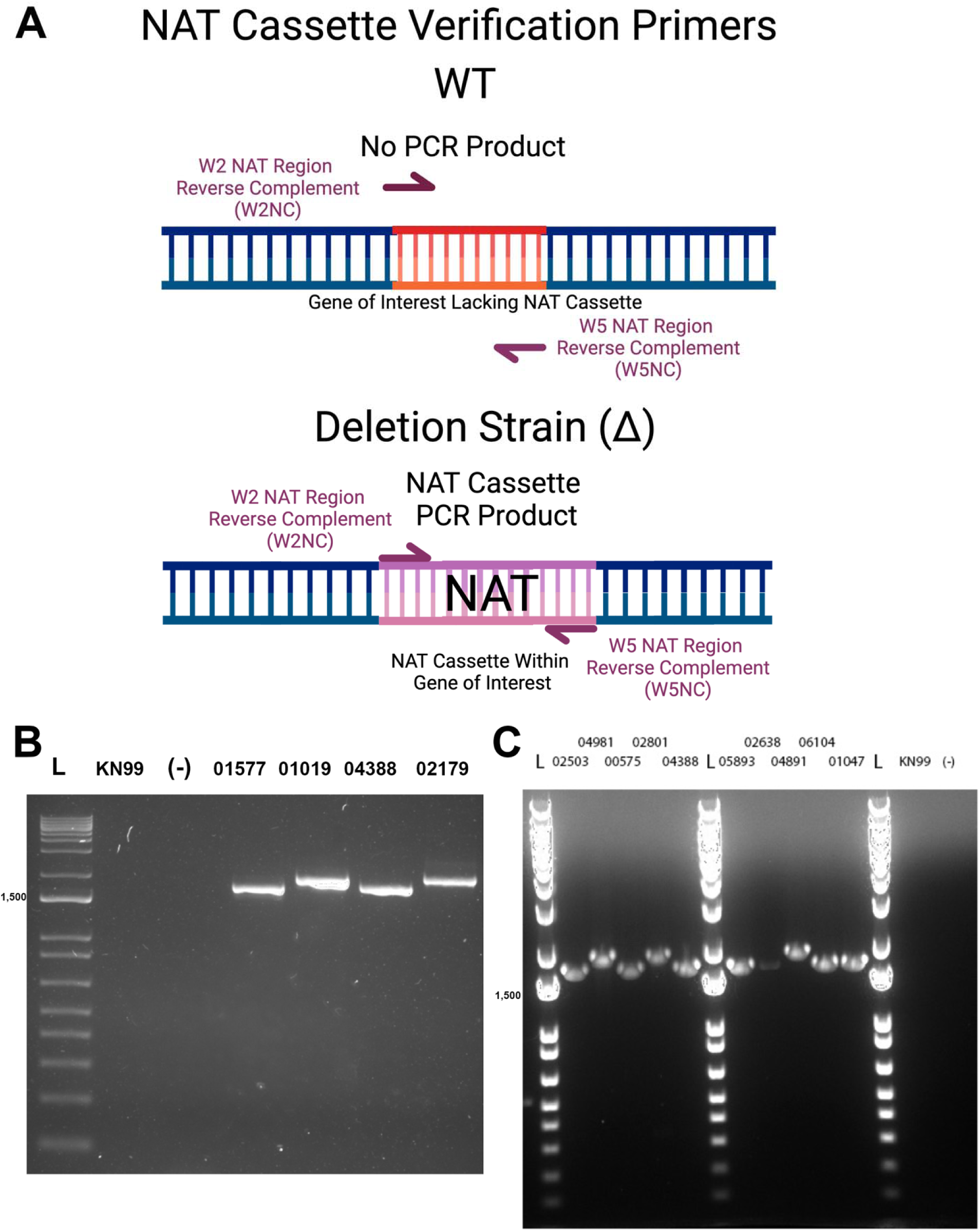
Comparing NAT cassette sizes between KO strains reveal size differences that correlate with locus-specific insertion failures **A)** NAT cassette verification methodology. The reverse complement of the NAT cassette-specific region of W2 (W2NC) and W5 (W5NC) were used to confirm the presence of the NAT cassette in putative KO strains. In WT KN99 , these primer pairs will not amplify a product due to a lack of insertion cassette. In KO strains, these primer pairs will only amplify the NAT cassette in any KO strain as there are no genomic-locus specific base pairs. **B)** NAT cassette size comparisons. KN99 and a no template control show lack of amplification. NAT cassettes PCR products from CNAG_02179Δ and CNAG_01019Δ, two strains that failed to have expected locus-specific disruptions, were larger in size than KO strains with correct KOs. **C)** Amplification of NAT cassette sequences by PCR across KO strains. This further confirmed the larger NAT cassette size correlated with locus-specific PCR failures. Note the larger NAT cassette in CNAG_04981Δ corresponds to the locus-specific PCR failure in figure 2 panel C. Both CNAG_01047Δ and CNAG_05893Δ contain smaller NAT cassettes and passed the PCR test from figure 2 panel C. We found the differences in NAT cassette size to be predictive of a KO strain’s locus-specific failure.

### RNAseq evidence to support a direct plate swap between the 2015 and 2008 KO collections

Following this hunch, we looked at the 2008 library plate map, available in the supplemental information of the accompanying paper [6]. We used data from an RNAseq experiment conducted on putative CNAG_02179Δ to confirm the plate swap (Figure 5A&B). CNAG_02179Δ was pulled from plate 2 well H12 from our 2015 labeled library plates. CNAG_03019Δ is the strain in plate 2 well H12 in the 2008 KO collection. RNAseq analysis provided definitive evidence demonstrating the presence of transcripts from the putative KO locus of CNAG_02179 and the absence of transcripts from the gene CNAG_03019, found in the analogous plate position in the 2008 collection (Fig 4A&B).

**Figure 4:**
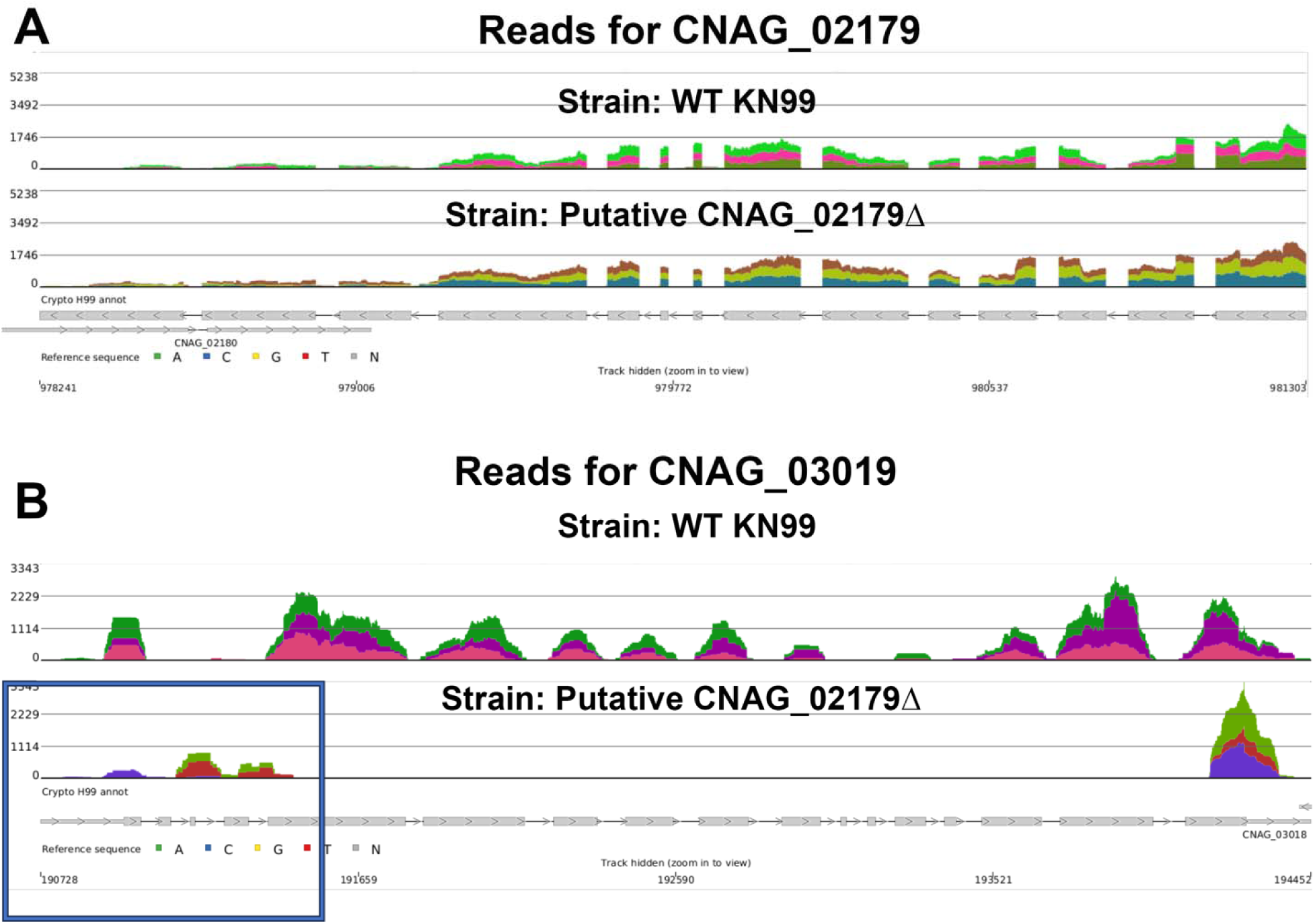
RNAseq reads for the putative CNAG_02179**Δ** strain revealed RNAseq reads were present for CNAG_02179 and absent at CNAG_03019 (the 2008 plate expected KO). **A)** RNAseq data comparing the coverage of CNAG_02179 in the WT KN99 strain (top) to the strain in our 2015 library plate 4 well H12 strain (putative CNAG_02179Δ) (bottom). Expression levels were identical in the WT and KO strain confirming no disruption in the expected locus. **B)** The RNAseq reads for CNAG_03019, the gene that would be knocked out if the plate was swapped directly with the 2008 library plate 4 well H12 strain in WT (top) and plate 4 well H12 strain (putative CNAG_02179Δ) confirming CNAG_03019 is the actual knocked out gene.

**Figure 5:**
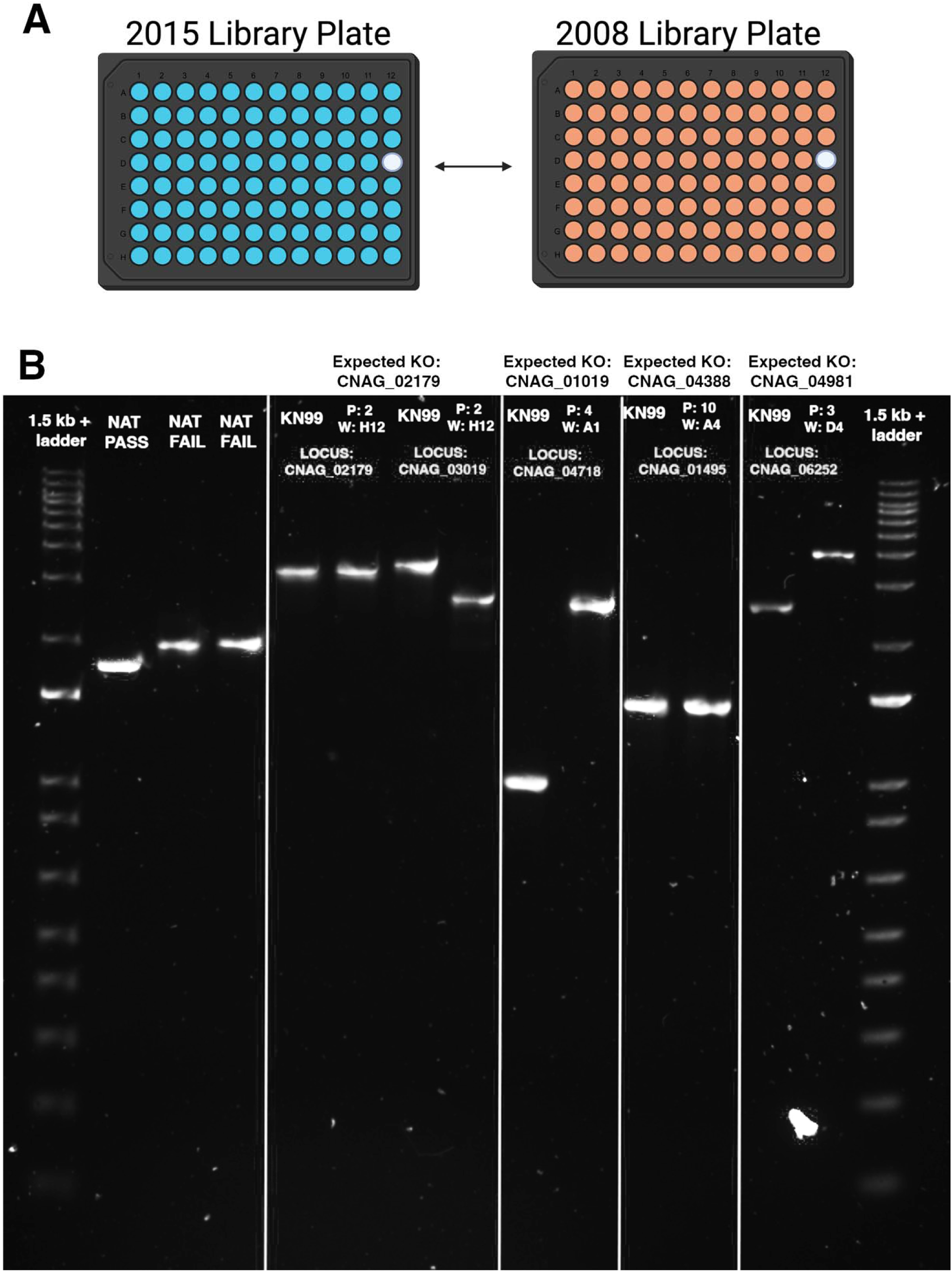
Summary PCR and confirmation of 2015 and 2008 direct plate swaps. **A)** Based on our findings whole 2015 plates were directly swapped with 2008 plates, not individual wells **B)** 1.5% agarose gel comparing PCR product sizes. L to R: 1 kb + ladder. The next three lanes are example NAT cassettes amplified using the strategy outlined in figure 3. The following lanes contain PCR products amplified using the locus-specific strategy outlined in figure 2. Expected KO indicates the CNAG that should be disrupted in the 2015 strain. Individual lane labels indicate genomic DNA source (KN99 WT or 2015 plate and well position of the strain). Locus labels indicate the genomic locus of the W2GC+W5GC primer pairs used for amplification of the 2008 swap loci. Of these strains, plate 2, plate 4, and plate 3 contained disruptions in the 2008 CNAG loci instead of the 2015 loci. The strain from plate 10 did not contain a disruption in the 2008 locus.

### PCR based evidence for a direct plate swap between the 2015 and 2008 KO collections

The evidence thus far suggested that some plates assembled into our copy of the 2015 KO collection were instead derived from another collection, most likely the earlier 14-plate collection on the avirulent H99W background from 2008 (Fig 5A). We tested this hypothesis using the W2GC + W5GC primer pairs for the expected genes if the 2015 plates were swapped with 2008 plates (Fig 5B). We tested this method on the KO strains from plate 2 well H12, plate 4 well A1, plate 10 well A4, and plate 3 well D4 (Fig 5B). Of these strains, plate 2, plate 4, and plate 3 contained disruptions in the 2008 CNAG loci instead of the 2015 loci. The strain from plate 10 did not contain a disruption in the 2008 locus, had a smaller NAT cassette, and the expected disruption in the 2015 locus.

Additionally, it is worth noting that the W2GC and W5GC primers designed from the 2015 sequences were used to amplify the suspected 2008 swapped gene disruptions (Fig 5A). The 2008 KO collection used different primers to KO target genes, therefore, can be expected to differ when comparing locus specific targets with 2015 library primers. For example, The PCR product amplified targeting the CNAG_03019 locus within the putative CNAG_02179Δ strain is larger than just the NAT cassette. This is explained by the insertion site of the 2008 version of this KO occurring ∼300 bp after the start codon. These findings are confirmed by the RNAseq data (Fig 4B). For locus-specific confirmation, 2008-specific primers could be used, but we feel the size differences add further evidence to confirm the KO is in fact of 2008 origin. Ultimately, we discovered that the larger NAT cassette size may be the result of a direct plate swap between the 2015 and 2008 KO collections.

### Our 2015 collection plates 1-7, 9, and 14 are 2008 KO collection plates

We tested many strains in our 2015 library using a mix of the genomic locus and NAT cassette size strategy (Figure 6A, Table 3). The most parsimonious interpretation of our results is that whole plates were directly swapped, not individual wells. The 2008 KO collection possesses 14 plates, and only plates within our first 14 plates of the 2015 KO collection were impacted. Plates #1-7, 9 and 14 of our 2015 deletion collection are instead KO strains from the 2008 collection, while plates #0, 8, 10-13, and 15-21 are the expected 2015 collection plates. The methodology comparing NAT cassette sizes between strains offers a quick, cost-effective methodology for testing the presence of a plate swap within a 2015 KO collection, as summarized in figure 7 (Fig 7A).

**Figure 6:**
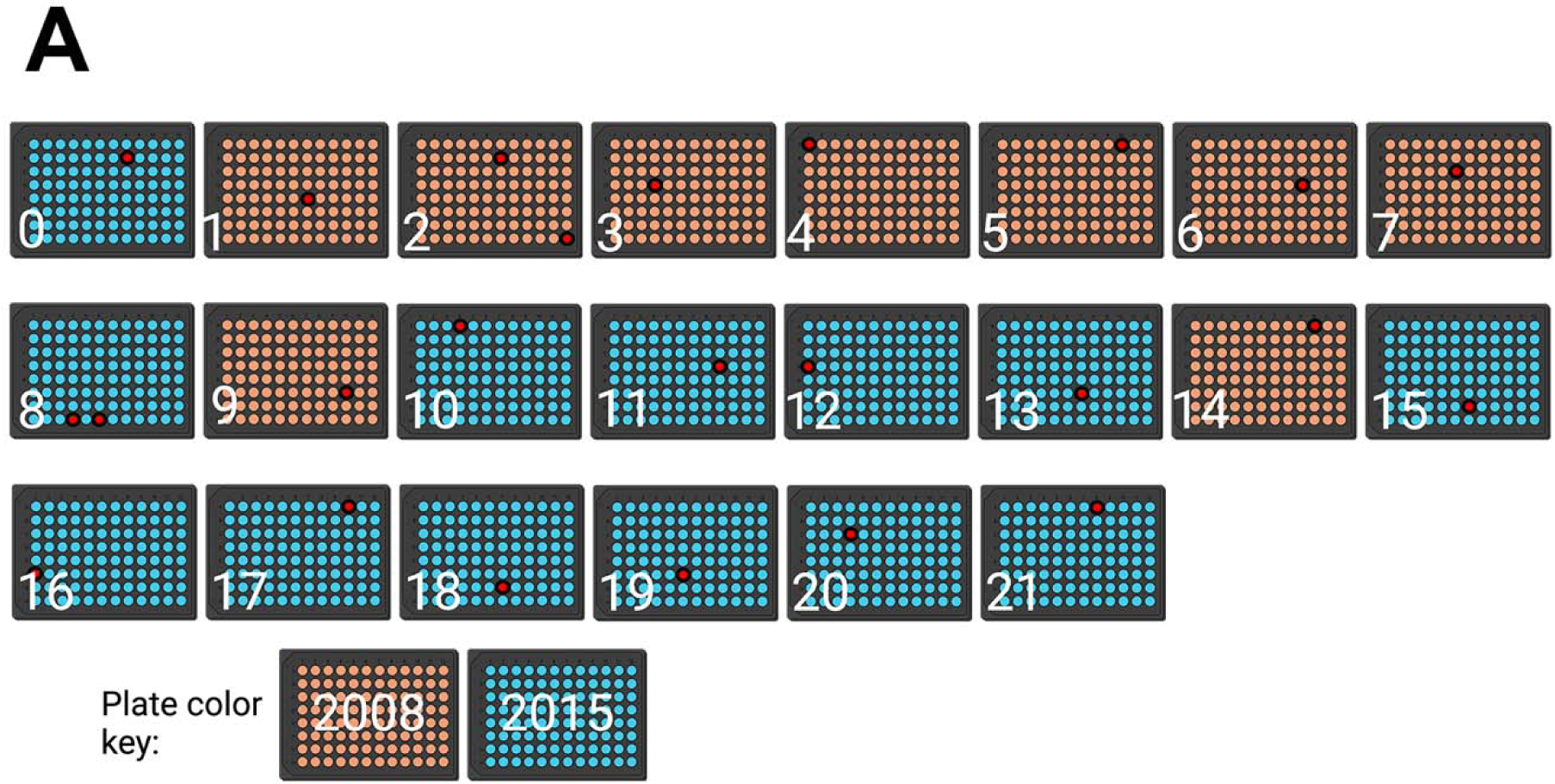
Plates 1-7, 9, and 14 are directly swapped **A)** Of the 22 plates in our 2015 KO collection, only the first 14 plates were impacted. Specifically, plates 1, 2, 3, 4, 5, 6, 7, 9, and 14 were directly swapped. Orange plates indicate 2008 swapped plates. Blue plates indicate 2015 library plates. Red wells indicate tested wells (Further information in Table 2)

**Figure 7:**
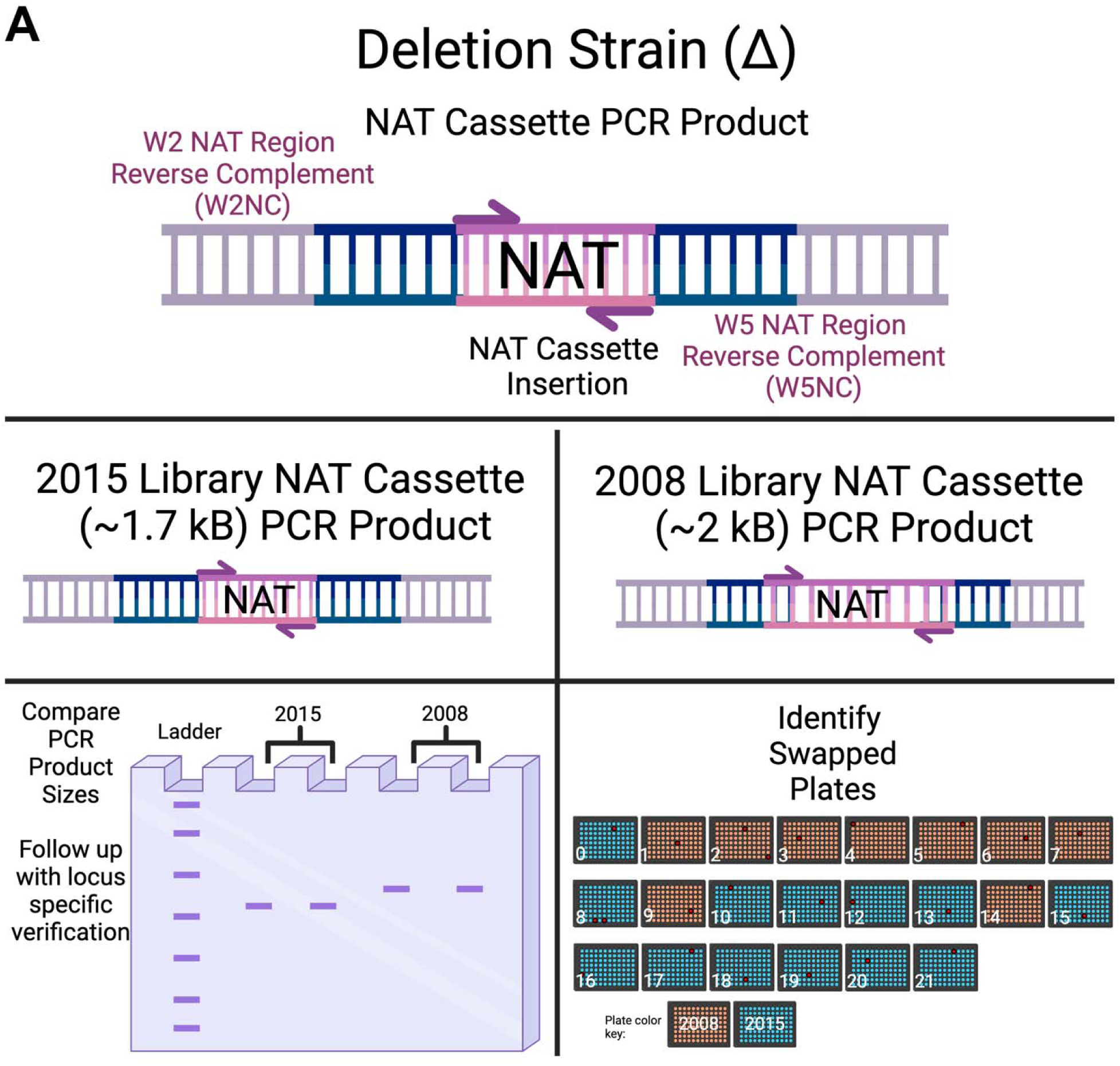
Graphical summary of library fidelity methodology **A)** In the process of uncovering the swapped plates within our 2015 KO collection, we found PCR amplifying 2008 library NAT cassettes and 2015 library NAT cassettes yielded different insertion sizes when compared by gel electrophoresis (∼2 kb vs ∼1.7 kb). By utilizing NAT cassette amplifying primers, strains can be compared regardless of genomic locus insertion site. This cost-effective simple method allows for comparisons across library plates to determine if a swap occurred using the same primers. Further verification using genomic DNA specific primers is necessary to fully confirm a swap.

## Discussion

Here, we describe an unfortunate plate mix-up involving cryptococcal gene KO libraries and demonstrate a cost-effective, time-saving method for authenticating the fidelity of *Cryptococcus neoformans* 2015 deletion KO collections. This will allow other labs to rapidly determine the status of their 2015 libraries. Using NAT cassette specific primers, not locus specific primers, to size comparison screening of the library proved a useful tool for initial screening. However, we did not test every strain and other sized NAT cassettes may exist within the library. Follow up with locus specific testing is essential to confirm that plates were swapped.

There are several limitations to the methods presented here. Firstly, we have found that locus specific primers were not specific enough to compare insertion sizes between 2008 and 2015 KO strains. The same insertion sites were not used between the 2008 and 2015 libraries to generate the same KO strains. Hence, care must be used when applying locus-specific primers to suspected 2008 swapped strains, as PCR product sizes will differ. An example of this can be seen when CNAG_03019Δ primers from the 2015 library are applied to the CNAG_03019Δ from the 2008 swapped plate. To confirm a 2008 swapped plate in a locus specific manner, we recommend using 2008 locus specific primers. Alternatively, be aware of the expected size differences when performing the experiment. Additionally, comparing NAT cassette sizes requires a 1.5% agarose gel because higher concentrations impede the separation of the PCR products. Finally, we have no information of where, when or how the plate swap occurred.

In summary, our experience is a cautionary tale of the foibles that can accompany the use of community resources like gene deletion libraries. Although these resources are powerful tools for discovery one must always maintain a certain skepticism when using them and validate any important finding by independent methods. We estimate that the lead authors in this paper lost approximately one year of research time in following up on preliminary data and then sorting out the mix-up with the libraries. Currently, these libraries are in many laboratories studying *C. neoformans* and continue to be useful tools for research. We are hopeful that our experiences and the validation roadmap described here will allow other laboratories to maximize their potential while avoiding false leads.

## Materials and methods

### Strains

Our lab purchased the 2015 and 2016 Madhani KO collections in March 2018 and the 2020 plate collections in March 2021.

### Genomic DNA isolation and colony PCR

Genomic DNA isolation was performed using the CTAB isolation method [13, 14]. Strains were cultured in 50 mL YPD, centrifuged and frozen overnight at 80 °C. The frozen cell pellet was powdered using glass beads and a bead beater. 10 mL CTAB extraction buffer (100 mM Tris pH7.5, 0.7 M NaCl, 10 mM EDTA, 1% CTAB, 1% beta-mercaptoethanol) was added to the culture pellet and vortexed. After 30 min of incubation at 65 °C, tubes were cooled in ice water and an equal volume of chloroform was added. The solution was mixed by inversion for 1 min and centrifuged for 10 min at max speed (RT). The top layer was transferred to a clean tube and an equal volume of isopropanol was added. The solution was mixed by inversion for 1 min and centrifuged for 5 min at max speed (RT). The supernatant was decanted off and 1 ml fresh 70% EtOH was added to the sample. After 5 min of centrifugation, the supernatant was decanted off and the pellet was resuspended in 100 µL sterile molecular biology water. For further testing of the entire library, we utilized colony PCR methodology [15]. In this method, small amounts of each strain are streaked directly into a PCR tube and microwaved for 2 min to release DNA. PCR mastermix and primers were added directly to the tube containing microwaved fungal cells.

Phusion PCR mastermix was used to amplify genomic loci and NAT cassettes. Due to the high annealing temperature of the NAT cassettes, a 2-step PCR protocol (combined annealing/extension) was utilized. All reactions were run on a 1.5% agarose gel. Higher percentages of agarose gels will impede size dependent comparisons.

### RNAseq

The yeasts were brought up to 1mL of Trizol and homogenized in the FastPrep 24 (MP Bio) with Lysing Matrix C Fast Prep tubes according to manufacturer’s protocol with minor modifications, followed by RNA extraction using the PureLink RNA Mini kit with on-column DNase treatment (ThermoFisher). RNA-Seq Libraries were prepared using the Universal Plus mRNA-Seq Library prep kit (Tecan Genomics) incorporating unique dual indexes. Libraries were assessed for quality by High Sensitivity D5000 ScreenTape on the 4200 TapeStation (Agilent Technologies). Quantification was performed with NuQuant reagent and by Qubit High Sensitivity DNA assay, on Qubit 4 and Qubit Flex Fluorometers (Tecan Genomics/ThermoFisher).

Libraries were diluted and an equimolar pool was prepared, according to manufacturer’s protocol for appropriate sequencer. An Illumina iSeq Sequencer with iSeq100 i1 reagent V2 300 cycle kit was used for the final quality assessment of the library pool. For deep mRNA sequencing, a 200 cycle (2 x 100 bp) Illumina NovaSeq 6000 S1 run was performed at Johns Hopkins Genomics, Genetic Resources Core Facility, RRID:SCR_018669. mRNA-seq data was analyzed with Partek Flow NGS Software as follows: pre-alignment QA/QC; alignment to *C. neoformans* Reference Index using STAR 2.7.8a; post- alignment QA/QC; quantification of gene counts to annotation model (Partek E/M); filter and normalization of gene counts; identification and comparison of differentially expressed genes with GSA (gene specific analysis). mRNA-seq data was aligned to *C. neoformans* using Bowtie 2 and chromosome view was used to visualize aligned reads to the putative CNAG_02179Δ strain vs WT KN99 strain (Partek Flow). Genome: NCBI: GCF_000149245.1_CNA3.

All sequence files and sample information have been deposited at NCBI Sequence Read Archive, NCBI BioProject: PRJNA1194623

## Acknowledgements

We would like to thank the Madhani Lab for their assistance in getting to the bottom of this issue and for providing the correct replacement strains to correct our papers.

